# HIV reverse transcriptase pre-steady-state kinetic analysis of chain terminators and translocation inhibitors reveals interactions between magnesium and nucleotide 3′-OH

**DOI:** 10.1101/2020.12.21.423855

**Authors:** Christopher R. Dilmore, Jeffrey J. DeStefano

## Abstract

Deoxythymidine triphosphate analogs with various 3′ sugar groups (-OH (dTTP), -H, -N_3_, -NH_2_, -F, -O-CH_3_, no group (2′,3′-didehydro-2′,3′-dideoxythymidine triphosphate (d4TTP)), and those retaining the 3′-OH but with 4′ additions (4′-C-methyl, 4′-C-ethyl) or sugar ring modifications (D-carba dTTP) were evaluated using pre-steady-state kinetics in low (0.5 mM) and high (6 mM) Mg^2+^ with HIV reverse transcriptase (RT). Analogs showed diminished incorporation rates (*k*) compared to dTTP ranging from about 2-fold (3′-H, -N_3_, and d4TTP with high Mg^2+^) to >10-fold (3′-NH_2_ and 3′-F with low Mg^2+^), while 3′-O-CH_3_ dTTP incorporated much slower than other analogs. Illustrating the importance of interactions between Mg^2+^ and the 3′-OH, *k* using 5 μM dTTP and 0.5 mM Mg^2+^ was only modestly slower (1.6-fold) than with 6 mM Mg^2+^, while analogs with 3′ alterations incorporated 2.8-5.1-fold slower in 0.5 mM Mg^2+^. In contrast, 4′-C-methyl and D-carb dTTP, which retain the 3′-OH, were not significantly affected by Mg^2+^. Consistent with these results, analogs with 3′ modifications were better inhibitors in 6 mM *vs*. 0.5 mM Mg^2+^. Equilibrium dissociation constant (K_d_) and *k*_pol_ determinations for dTTP and analogs lacking a 3′-OH indicated that low Mg^2+^ caused a several-fold greater reduction in *k*_pol_ with the analogs but had little effect on K_d_, results consistent with a role for 3′-OH/Mg^2+^ interactions in catalysis rather than nucleotide binding. Overall, results emphasize the importance previously unreported interactions between Mg^2+^ and the 3′-OH of the incoming nucleotide and suggest inhibitors with 3′-OH groups may have advantages in the low free Mg^2+^ in physiological settings.

## Introduction

Nucleoside Reverse Transcriptase Inhibitors (NRTIs) are a hallmark of Antiretroviral Therapy (ART). All NRTIs currently approved for ART are “chain terminators” and either lack a hydroxyl group at the 3′ ribose position or have a substituted sugar or other group replacing ribose. In all cases there is no group present for addition of the next base by reverse transcriptase (RT) (reviewed in: (1–3)). A second class of emerging NRTIs referred to as “translocation inhibitors” retain the 3′-OH but contain modifications at other positions that dramatically slow the addition of the next base (4–7). The first approved HIV drug, azidothymidine (also referred to as Zidovudine, AZT, or ZDV), is essentially dTTP with the 3′-OH replaced by an azido group. Dideoxy drugs replace the 3′-OH with H (including dideoxyinosine (also referred to as Didanosine or ddI) and dideoxycytidine (a formerly approved HIV drug also referred to as Zalcitabine or ddC). Other drugs remove both the 3′-OH and a hydrogen atom from the 3′ position by addition of a carbon double bond to the ribose ring (2′,3′-Didehydro-3′-deoxythimidine (also referred to as Stavudine or d4T) and the guanosine analog Abacavir (ABC)). Finally, some approved drugs substantially modify or remove the ribose sugar including Lamivudine (3TC), Emtricitabine (FTC), and Tenofovir (TDF). Despite the success of drugs lacking a 3′-OH, for some translocation inhibitors this group is pivotal in promoting the phosphorylation of the prodrug in cells (8).

Others have examined the effect of different groups at the 3′ position of thymidine and adenosine analogs for inhibition of HIV RT (9, 10). For thymidine, inhibition assays using steady-state kinetics and measuring incorporation on poly(rA)-oligo(dT_10_) indicated that substitutions of fluorine (F), amino (NH_2_), azido (N_3_), and hydrogen (H) at the 3′ position were all effective inhibitors of dTTP incorporation with modest overall differences, with 3′-F and 3′-N_3_ being the most and least effective, respectively (9). Despite this, only the 3′-N_3_ derivative (marketed as AZT) has been an effective HIV drug. Drug effectiveness is complicated by several factors including stability, uptake, cellular phosphorylation, target and off-target utilization, and resistance profiles among others. Even small modification could potentially affect any of these steps. The F and H 3′ substitution (as well as several other changes at other positions) on dATP were also tested for anti-HIV activity and cell toxicity (10). Both analogs had lower cytotoxicity than AZT, however, they also had much lower potency. Still, the selectivity index for 3′-H (ddATP) was only about 4.5-fold lower than AZT suggesting it could be an effective drug.

Previous results indicated that the level of Mg^2+^ in *in vitro* reactions can affect the fidelity of HIV RT, efficiency of reverse transcription, and the potency of both NRTIs and Nonnucleoside Reverse Transcriptase Inhibitors (NNRTIs). Fidelity increased significantly (11, 12) in reactions performed in low Mg^2+^ (0.25-0.5 mM) that more closely mimics cellular free Mg^2+^ conditions (13–17), in comparison to the high Mg^2+^ (~6 mM) typically used in *in vitro* reactions with RT. The efficiency of reverse transcription, as judged by the length of products produced in primer extension reactions, was greater with physiological Mg^2+^ concentrations (18). The effect on NRTIs and NNRTIs was opposite, with the former being less effective in low Mg^2+^ and the latter being more effective (18, 19).

In this report we used pre-steady-state kinetic analysis at a fixed nucleotide concentration (5 μM) that approximated cellular dNTP concentrations (20–22), and high (6 mM) or low (0.5 mM) Mg^2+^ concentrations to examine the incorporation rate of dTTP and several dTTP analogs with substitutions at the 3′ position. This included 3′-OH (dTTP), -N_3_ (AZT), -NH_2_, -F, -O-CH_3_, -H (ddTTP), and d4T (no group at this position). Three translocation type dTTP analog inhibitors (4′-C-methyl, 4′-C-ethyl, and D-carba dTTP) that retained the 3′-OH were also tested. Determinations of *k*_cat_ and K_d_ were also performed for some analogs under pre-steady-state conditions. Overall, the results illustrate a potential benefit for NRTI drugs with 3′-OH groups, especially under physiological Mg^2+^ concentrations, while also uncovering a previously unreported direct or indirect interaction between the 3′-OH of incoming nucleotides and Mg^2+^.

## Results

### Substitution of the 3′-OH differentially affects the incorporation rate of dTTP analogs

All the analogs tested, including chain terminators and translocation inhibitors, were incorporated more slowly than dTTP in both 0.5 and 6 mM Mg^2+^, in pre-steady-state assays with 5 μM nucleotide (Table 1, see Fig. 1 for an example of an analysis). This amount was chosen as it approximately mimics the level of nucleotides in dividing T cells (20–22). The incorporation rate ordered from highest to lowest in 6 mM Mg^2+^ was: 3′-OH > (-N_3_, no group (d4TTP), -H) > (4′-C-methyl, 4′-C-ethyl, -NH_2_) > (-F, D-carba) ≫ -O-CH_3_. In 0.5 mM Mg^2+^ the order was: 3′-OH > (4′-C-methyl) > (-N_3_, no group, -H, 4′-C-ethyl, D-carba, -NH_2_) > -F ≫ -O-CH_3_. The 3′-O-CH_3_ analog was by far the slowest and no data could be obtained using pre-steady-state conditions. However, steady-state analysis showed that the analog could be incorporated by RT (Fig. 2). Among the chain terminating analogs, at 6 mM Mg^2+^, an -N_3_, -H, or no group at the 3′ position were most tolerated. The 3′-F was modestly slower than 3′-NH_2_ and the incorporation rate of 3′-O-CH_3_ was negligible in comparison (see above). Note that these results agree with a previous steady-state analysis of some of these analogs indicating that they were efficiently incorporated by HIV RT ((9) and Introduction). However, the relative potency of the various inhibitors did not agree with our pre-steady-state analysis.

**Table 1.**
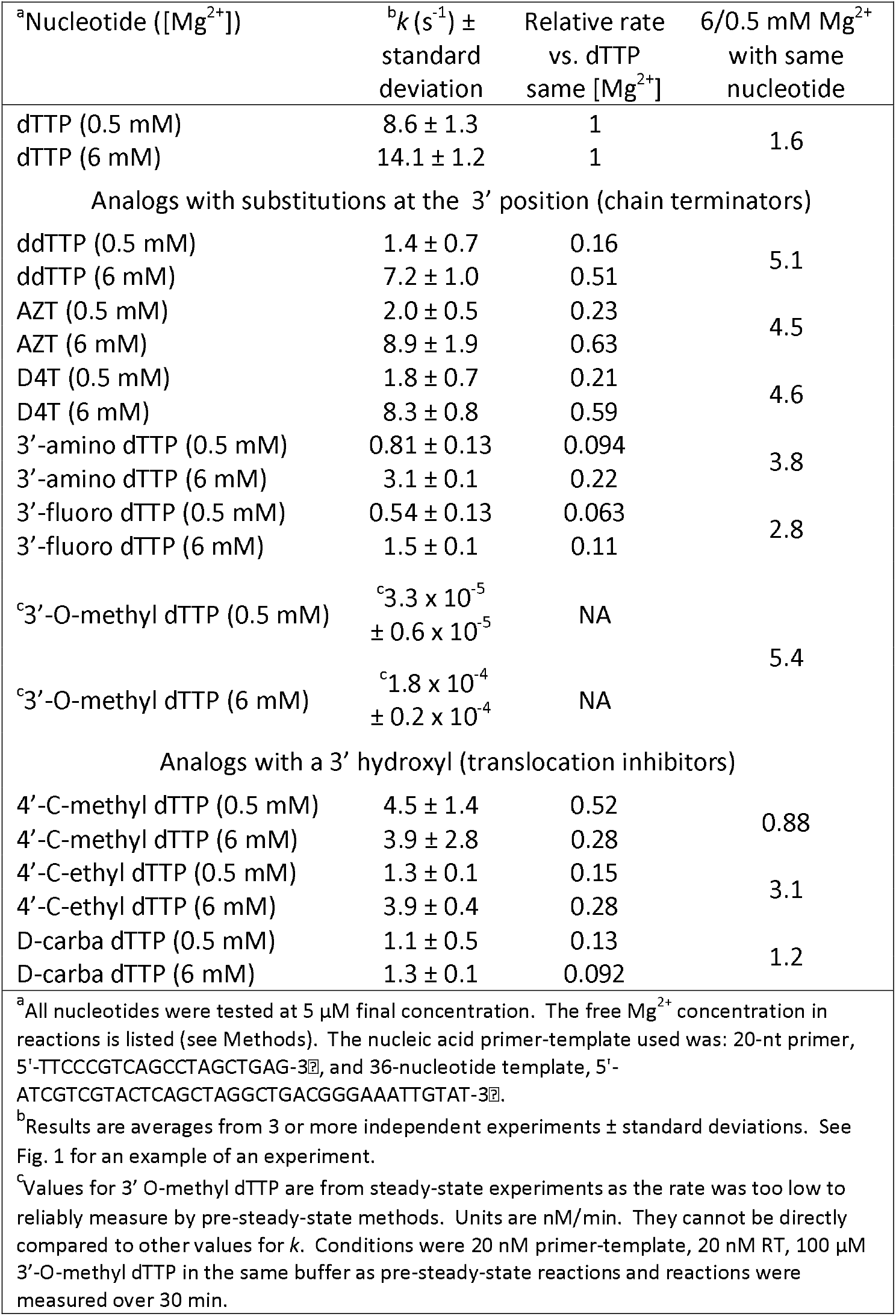
Comparison of pre-steady state incorporation of dTTP analogs.

**Figure 1.**
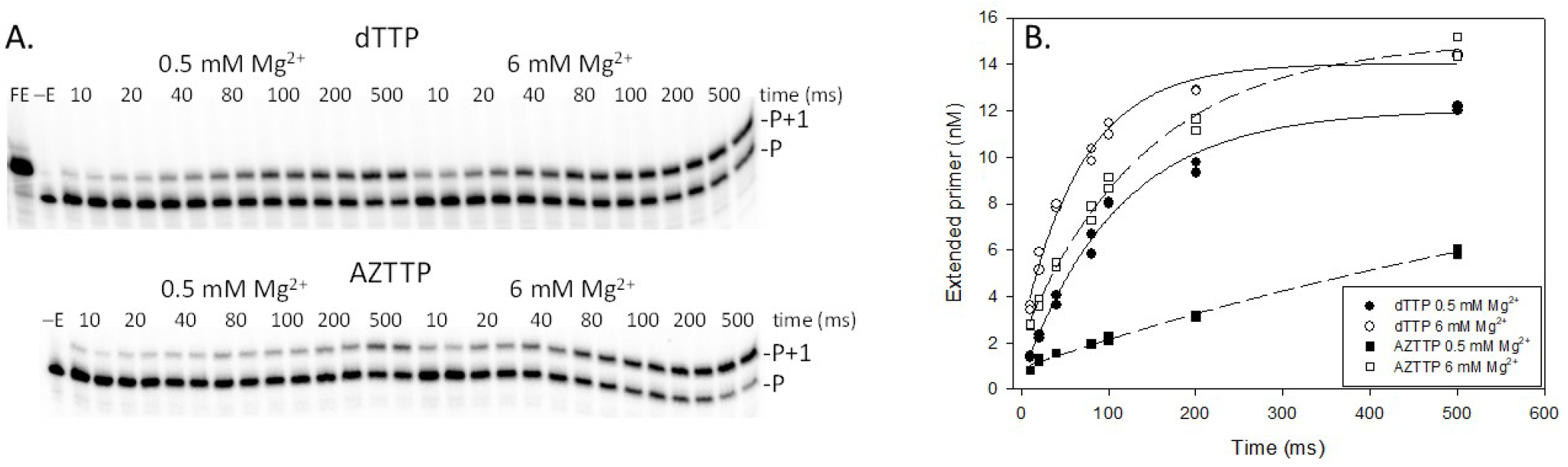
Example of pre-steady state determination of the rate (*k*) of nucleotide addition with 5 μM dTTP or AZTTP and 0.5 or 6 mM Mg^2+^. (A) Assays were conducted as described in Materials and Methods with a 20-nt 5′-^32^P end-labeled DNA primer bound to a 36-nt DNA template using rapid quench analysis. Time points in these assays were 10, 20, 40, 80, 100, 200, and 500 ms and were conducted in duplicate. For nucleotide analogs with slower incorporation rates longer time points were also used. Positions of the primer (P) and primer + 1 nucleotide (P+1) are shown for assays with dTTP and AZTTP at 0.5 or 6 mM Mg^2+^. FE, full extension; -E, no enzyme added. (B) The data was fitted to an exponential equation as described in Materials and Methods to yield the incorporation rate (*k (s^−1^)*) used to produce the data in Table 1.

**Figure 2.**
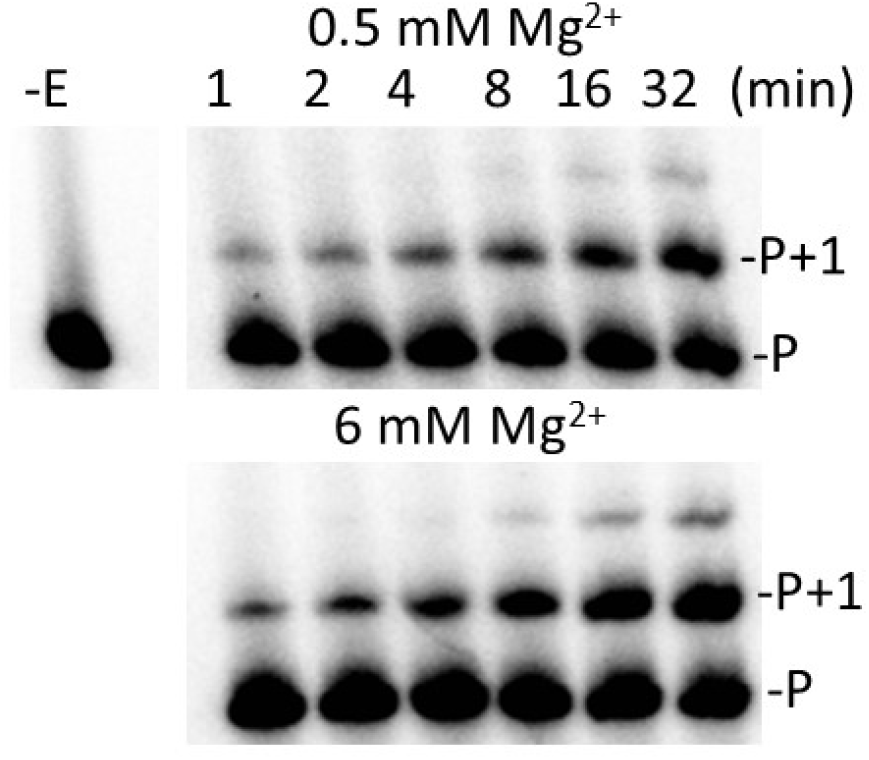
Addition of 3′-O-CH_3_ dTTP in steady-state assays with HIV RT with 0.5 or 6 mM Mg^2+^. Reactions were carried out as described in Materials and Methods using a 20-nt 5′-^32^P end-labeled DNA primer bound to a 36-nt DNA template. All reactions contained 100 μM 3′-O-CH_3_ dTTP with either 0.5 or 6 mM Mg^2+^. Reactions were carried out for the indicated times and resolved on a 20% denaturing gel. The proportion of extended primer was quantified on a phosphorimager and used to determine the rate (*k*) of nucleotide addition. -E, no enzyme added; P, 5′-^32^P end-labeled DNA primer; P+1, primer extended with 3′-O-CH_3_ dTTP.

Incorporation rates for translocation inhibitors were less dependent on the Mg^2+^ conditions. Rates for the 4′-C-methyl and 4′-C-ethyl derivatives were similar in 6 mM Mg^2+^ and both were about 3-fold faster than D-carba dTTP. In 0.5 mM Mg^2+^, only the incorporation rate for 4′-C-ethyl dTTP declined significantly.

### The effect of lower Mg^2+^ on incorporation is largely independent of the type of 3′ substitution

All analogs that did not have a 3′-OH were incorporated more slowly in 0.5 than 6 mM Mg^2+^. The magnitudes of the decreases were in a range between 2.8-5.1-fold (Table 1), with 3′-F and 3′-H (ddTTP) being the lowest and highest, respectively. However, the difference between the various 3′ analogs was small and, in most cases, not significant. This indicates that the low Mg^2+^ effect results more from the loss of the 3′-OH than a property of the substitution. The 4′-C-methyl and D-carba translocation inhibitor analogs were not strongly affected by lower Mg^2+^ while dTTP showed just a 1.6-fold decrease in incorporation rate with low Mg^2+^ (Table 1). Curiously, 4′-C-ethyl dTTP behaved more like the chain terminators and was incorporated about 3 times more slowly in 0.5 mM Mg^2+^. It is notable, however, that despite possessing a 3′-OH, 4′-C-ethyl dTTP actually behaves like a chain terminator as unlike 4′-C-methyl and D-carb dTTP, nucleotide cannot be added to this moiety ((23) and see Discussion).

### Analogs without a 3′-OH demonstrate a more profound loss of catalytic efficiency in lower Mg^2+^

The observed incorporation rate of a nucleotide or analog at a particular concentration is dependent on its affinity for enzyme as well as the overall rate of the catalytic steps. Several reports have shown that 3′-OH substitutions may alter both the binding affinity (i.e. equilibrium dissociation constant (K_d_)) for HIV RT as well as the maximum incorporation rate (*k*_pol_) (29–33). We choose three dNTPs/analogs, dTTP, AZTTP, and d4TTP, to evaluate more thoroughly and determine both pre-steady-state K_d_ and *k*_pol_ values at 0.5 and 6 mM Mg^2+^. Although it would have been more complete to test all 10 analogs, a combination of limiting amounts of material and time made it more feasible to test a smaller representative set. The decision to focus on AZTTP and d4TTP was made because there is extensive literature on these compounds, and they are approved HIV drugs. The K_d_ values for all 3 compounds were similar with d4TTP showing a modestly higher K_d_ than the others at both Mg^2+^ concentrations (Fig. 3 and Table 2). The level of Mg^2+^ did not strongly affect the K_d_ value for any of the nucleotides. Values for K_d_ at 6 mM obtained hear were comparable to previous results with high Mg^2+^ (33). As expected from the results in Table 1, changes in *k*_pol_ were more pronounced, especially for the nucleotides lacking a 3′-OH (AZTTP and d4TTP). The decline in catalytic efficiency (*k*_pol_/K_d_), with 0.5 vs. 6 mM Mg^2+^ was also more pronounced for these analogs. For example, AZTTP was incorporated only 1.5-fold less efficiently than dTTP in 6 mM Mg^2+^, but 7.2-fold less efficiently in 0.5 mM Mg^2+^ (Table 2). The data again illustrates that the lack of a 3′-OH has a more pronounced negative affect on incorporation in low, more physiological Mg^2+^, and a decrease in incorporation rate (*k*) as opposed to changes in affinity (K_d_) for the analogs is the predominant factor.

**Table 2.**
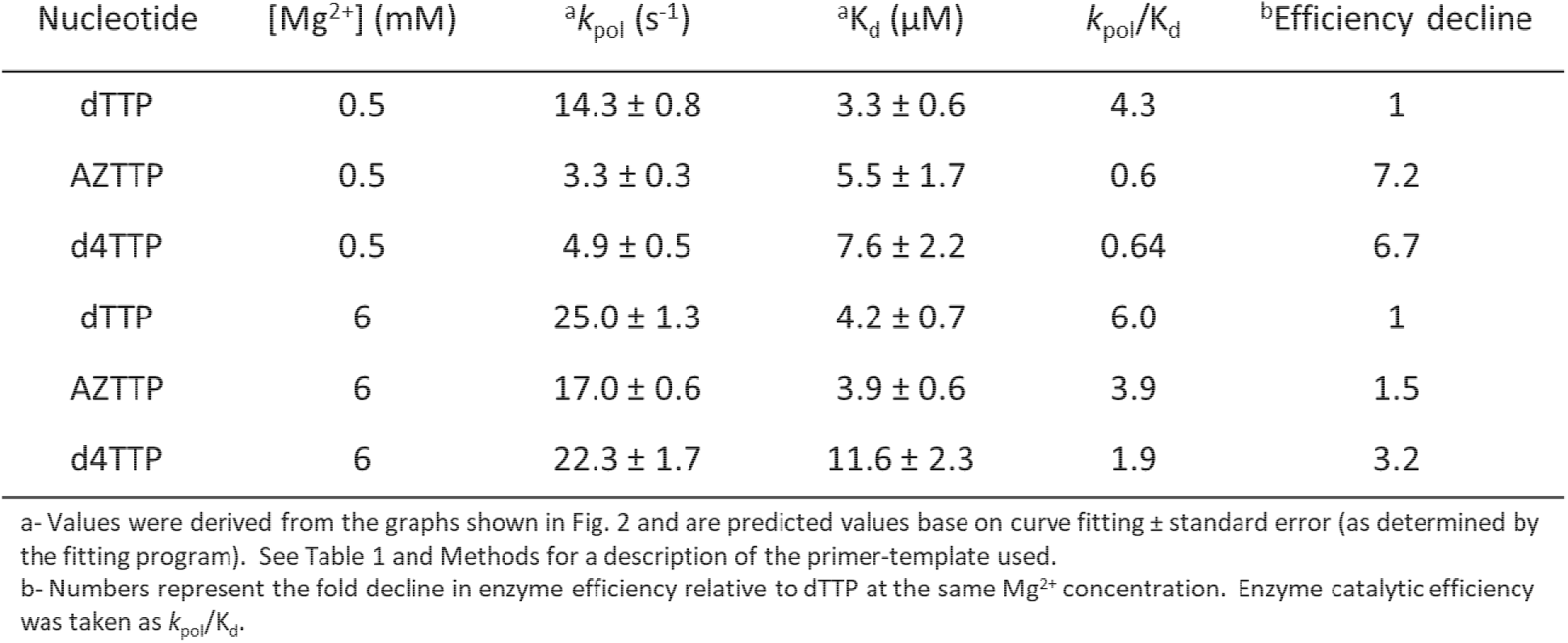
Pre-steady state kinetic data for HIV RT incorporation of TIP, AZTTP, and D4TTP.

| Nucleotide | [Mg<sup>2+</sup>] (mM) | <sup>a</sup>$k_{\text{pol}}$ (s<sup>-1</sup>) | <sup>a</sup>$K_{\text{d}}$ (μM) | $k_{\text{pol}}/K_{\text{d}}$ | <sup>b</sup>Efficiency decline |
| --- | --- | --- | --- | --- | --- |
| dTTP | 0.5 | 14.3 ± 0.8 | 3.3 ± 0.6 | 4.3 | 1 |
| AZTTP | 0.5 | 3.3 ± 0.3 | 5.5 ± 1.7 | 0.6 | 7.2 |
| d4TTP | 0.5 | 4.9 ± 0.5 | 7.6 ± 2.2 | 0.64 | 6.7 |
| dTTP | 6 | 25.0 ± 1.3 | 4.2 ± 0.7 | 6.0 | 1 |
| AZTTP | 6 | 17.0 ± 0.6 | 3.9 ± 0.6 | 3.9 | 1.5 |
| d4TTP | 6 | 22.3 ± 1.7 | 11.6 ± 2.3 | 1.9 | 3.2 |

b- Numbers represent the fold decline in enzyme efficiency relative to dTTP at the same Mg<sup>2+</sup> concentration. Enzyme catalytic efficiency was taken as $k_{\text{pol}}/K_{\text{d}}$ .

**Figure 3.**
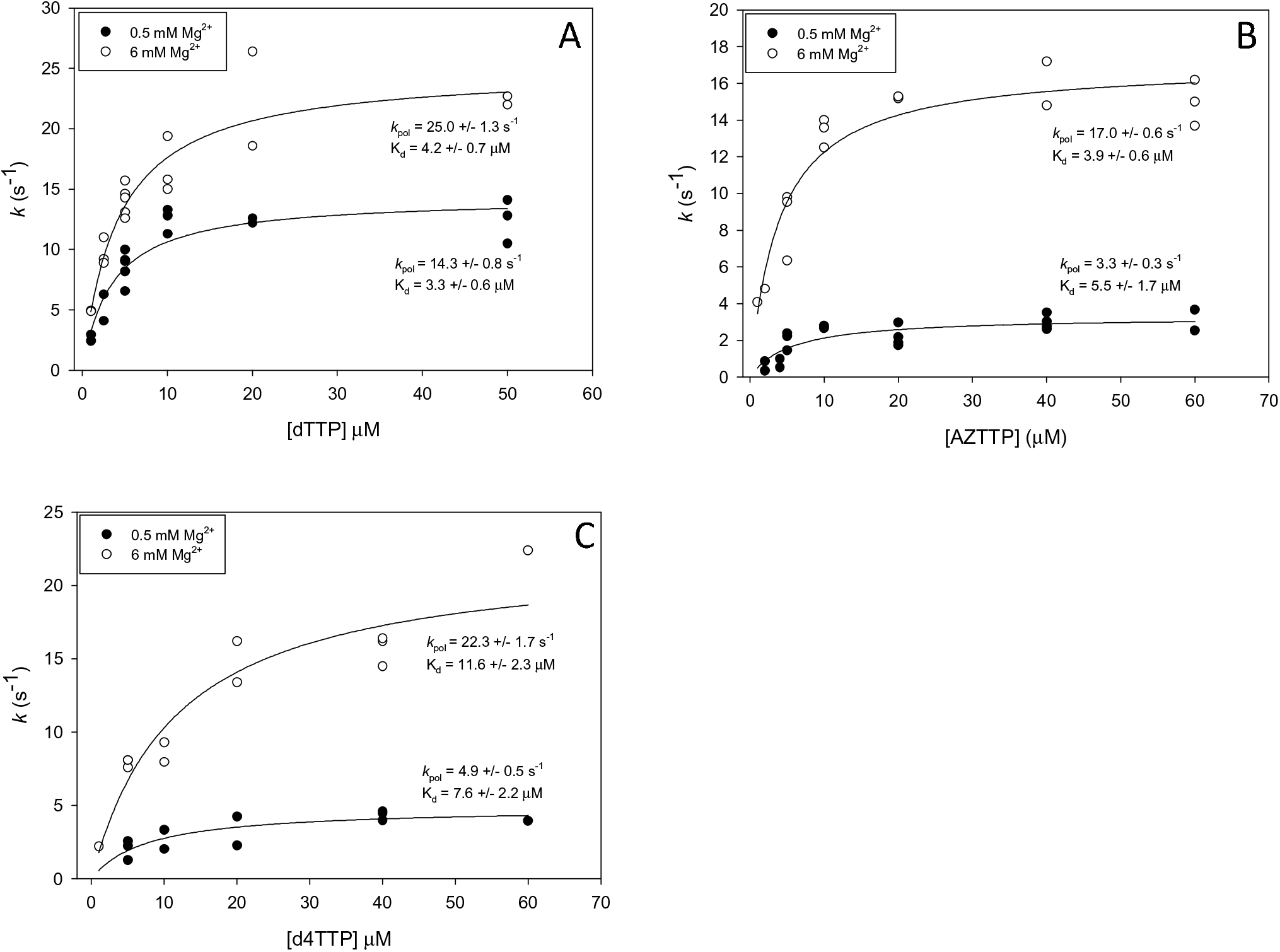
Pre-steady state determination of equilibrium dissociation constant (K_d_) and *k*_pol_ for dTTP, AZTTP, and d4TTP. Assays were conducted as described in Materials and Methods with a 20-nt 5′-^32^P end-labeled DNA primer bound to a 36-nt DNA template using rapid quench analysis. Data points are from independent experiments at various concentrations of dTTP (A), AZTTP (B), or d4TTP (C) (see Fig. 1 for an example). Points from several experiments (typically 2-3, while only 1 point was used for the highest d4TTP concentration), were plotted and fit to an equation to determine values as described in Materials and Methods. The indicated values for K_d_ and *k_pol_* are derived from the curve fit program. Values are ± standard errors as determined by the program.

### Inhibition assays on a heteropolymeric DNA template are generally consistent with the kinetic analysis

To evaluate the effectiveness of the various analogs at terminating DNA synthesis, a primer extension assay was performed on a 100-nt DNA template primed with a 20-nt 5′-^32^P labeled DNA primer (Fig. 4). Assays contained 5 μM dCTP, dGTP, dATP, and dTTP. Analogs were included at 3 μM when present. Consistent with 6/0.5 mM Mg^2+^ ratios shown in Table 1, all chain terminators and 4′-C-ethyl dTTP were less effective (as judged by a lower level of paused (terminated) products and an increase in fully extended products) in 0.5 mM Mg^2+^ compared to 6 mM Mg^2+^. Among translocation inhibitors, 4′-C-methyl dTTP was essentially equivalent at both 0.5 and 6 mM Mg^2+^ while D-carba was a weak inhibitor under both conditions. The 3′-F analog was the weakest inhibitor among the chain terminators and was also incorporated the slowest in pre-steady-state reactions (Table 1, note that 3′-O-CH_3_ produced no inhibition in this assay (data not shown) and its incorporation was not measurable in pre-steady-state reactions (see above)). However, it is important to note that the inhibition observed in these reactions is a function of both incorporation kinetics and binding affinity of the analog for RT vs. binding affinity for dTTP. Therefore, a direct correlation between the results in Table 1 and those in the inhibition assay would not necessarily be expected. Finally, results in Table 1 were performed using a single sequence position on a primer-template while the experiments in Fig. 4 use a long template with positions for thymidine incorporation presented in several sequence contexts.

**Figure 4.**
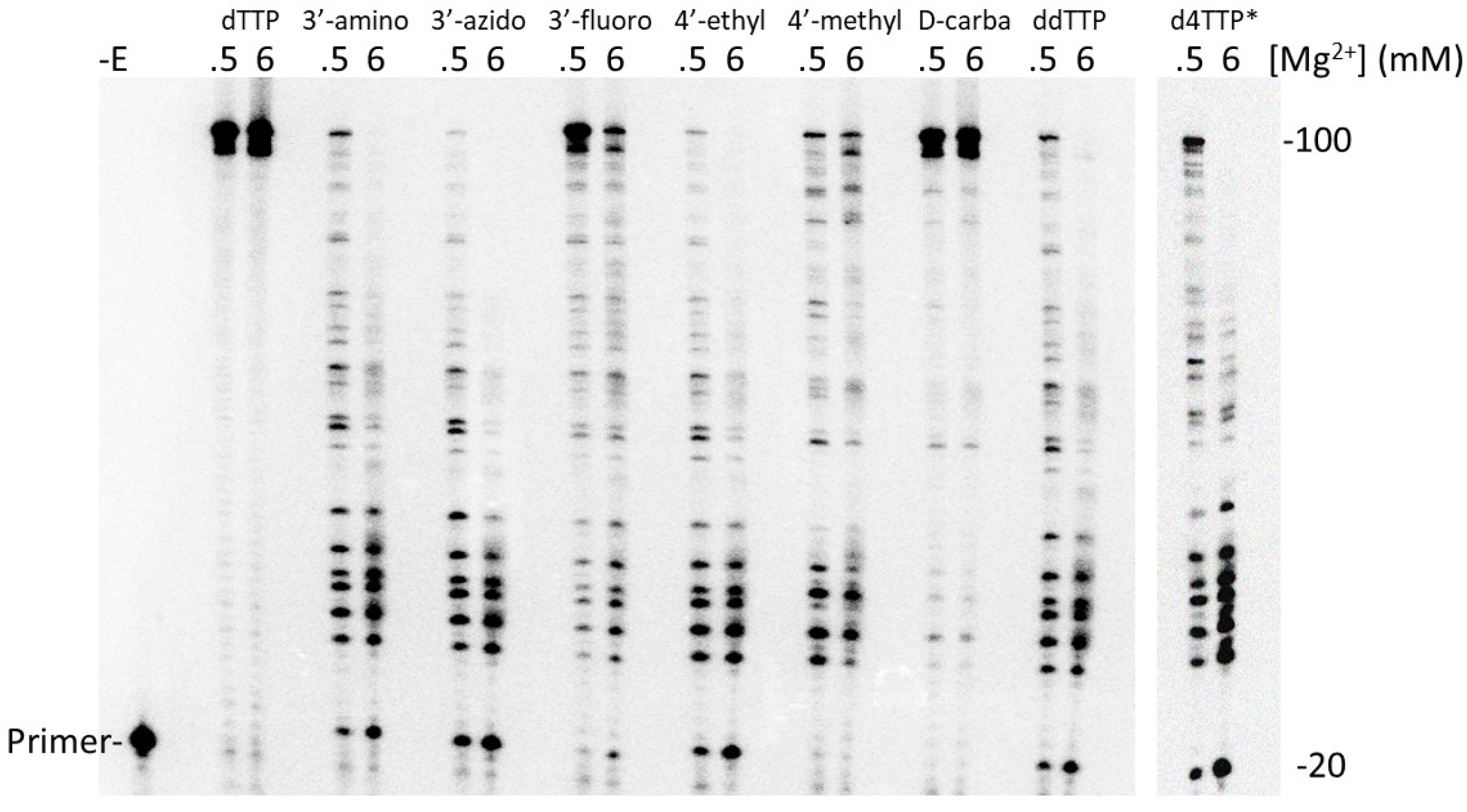
Primer extension on a 100-nt DNA template with HIV RT. Reactions were carried out as described in Materials and Methods using a 20-nt 5′-^32^P end-labeled DNA primer bound to a 100-nt DNA template. All reactions contained 5 μM dNTPs and 3 μM of the indicated dTTP analog with either 0.5 or 6 mM Mg^2+^. Reactions were carried out for 30 min and resolved on an 8% denaturing gel. -E, no enzyme added. *d4TTP lane came from another gel and a separate experiment.

## Discussion

In this report, several chain-terminating 3′-OH substitutions of dTTP were tested for incorporation by HIV RT. All the analogs with 3′ alterations were incorporated and several (e.g. AZT, ddTTP, and d4TTP) with only modestly slower rates (~2-fold) than dTTP in 6 mM Mg^2+^ (Table 1). In contrast, the rate of incorporation compared to dTTP dropped to ~4.5-16-fold slower (excluding 3′-O-CH_3_ which was much slower at both high and low Mg^2+^) in 0.5 mM Mg^2+^. This is consistent with previous experiments showing that NRTIs are better inhibitors with higher Mg^2+^ concentrations *in vitro* (19), an observation that was also shown for analogs tested here (Fig. 4). Overall, the results support a previously unreported interaction, either direct or indirect, between Mg^2+^ and the 3′-OH of incoming nucleotides. Further, the results suggest that this interaction plays a role in catalysis rather than nucleotide binding affinity (see below).

The importance of the 3′-OH was supported by experiments with translocation type inhibitors (4′-C-methyl and D-carba dTTP) which retain the 3′-OH and behaved more like dTTP in that their rate of incorporation was less affected by low Mg^2+^ (Table 1). However, 4′-C-ethyl dTTP was an exception, showing a 3.1-fold lower rate of incorporation in low Mg^2+^ compared to high. As was noted, this analog terminates further extension by RT, in contrast to 4′-C-methyl and D-carba dTTP which delay extension (23). This suggests that the 3′-OH may be in the wrong position for catalysis and this position may also weaken the proposed interactions of the 3′-OH with Mg^2+^. The translocation inhibitor 4′-ethynyl-2-fluoro-2′-deoxyadenosine triphosphate (EFdATP) is currently in clinical trials. It also binds strongly to HIV RT as the ethynyl group can access a hydrophobic pocket in RT which may stabilize binding (7, 34). This, and favorable kinetics lead to EFdATP showing a competitive advantage over the natural dATP nucleotide (35). EFdATP inhibition is also insensitive to Mg^2+^ concentration (19), and incorporated EFdA can accept additional nucleotides at a highly reduced rate (35). This suggests that translocation inhibitors with the 3′-OH positioned for nucleotide addition behave more like natural nucleotides with respect to Mg^2+^ interactions.

It was interesting the 4′-C-ethyl dTTP was clearly a more effective inhibitor than 4′ C-methyl or D-carba dTTP at 6 mM Mg^2+^ during extension on the long template (Fig. 4). Even at 0.5 mM Mg^2+^, where 4′-C-ethyl dTTP was incorporated much more slowly than 4′-C-methyl dTTP and at a rate similar to D-carba dTTP (Table 1), it still was modestly more effective than 4′-C-methyl dTTP and much more effective than D-carba dTTP in the template inhibition assay. This may be due to two factors. First, 4′-C-ethyl dTTP, as stated above, behaves like a chain terminator, and does not allow the addition of other nucleotides after incorporation (23). Additional nucleotides can be added with delayed kinetics with 4′-C-methyl and D-carba dTTP (23, 24) and, for these analogs, extended products in the reactions in Fig. 4 may include DNA chains containing more than one inhibitor nucleoside. Second, the K_d_ for 4′-C-ethyl, 4′-C methyl and D-carba dTTP binding to HIV RT was measured at 6 mM Mg^2+^ in a separate analysis on a different template (manuscript in preparation). The K_d_ value for D-carba dTTP was ~4-8-fold higher than the others. This helps explain why D-carba dTTP is by far the weakest inhibitor among both translocation and chain terminating inhibitors in the template extension inhibition assay (Fig. 4). It binds more weakly to RT than the other translocation inhibitors, and this, coupled with a relatively low incorporation rate (Table 1), weakens inhibition.

Although positioning of the 3′-OH of an incoming nucleotide is coordinated through many interactions with HIV RT, glutamine 151 (Q151) presumably interacts directly with this group through a proposed hydrogen bond between the amide oxygen from Q151 and the 3′-OH hydrogen atom. Mutations in Q151 like Q151N (glutamine to asparagine) that disrupt this interaction weaken dNTP binding but do not significantly affect *k*_pol_ (29). This indicates that Q151 likely plays a key role in stabilizing dNTP binding. Dideoxy dTTP which lacks a 3′-OH demonstrated weaker binding to HIV RT than dTTP. However, binding relative to dTTP improves with Q151N which has weakened association with the 3′-OH. This again predicts a role for Q151 in stabilizing binding of dNTPs through interactions with the 3′-OH, as the advantage of a 3′-OH for binding is mitigated when Q151 is altered to a non-interacting amino acid.

High divalent cation concentrations can also improve incorporation of analogs without 3′-OH groups with other polymerases, similar to what was found here. The Klenow fragment of *E. coli* polymerase I (KF) discriminates strongly against ddNTPs relative to dNTPs. However, ddTTP incorporation improves in high Mg^2+^ showing an optimum of 25 mM which is several-fold greater than the optimal concentration for incorporation of dTTP (~2 mM) (36). Like Q151N for HIV RT, an E710A (glutamic acid to alanine) KF mutation reduced ddNTP/dNTP discrimination, consistent with a role for this amino acid in interactions with the 3′-OH. The authors hypothesize that Mg^2+^ may bridge an interaction between KF E710 and the 3′-OH group of incoming nucleotides, although more complex explanations where E710 interacts with Mg^2+^ *via* repositioning of other active site residues could not be ruled out by the data. It was also not clear if the Mg^2+^ ion in question was metal ion A and/or B, the putative metal ions involved in nucleotide catalysis at polymerase active sites, or another, as yet undescribed metal ion. In this regard, it is interesting that a third metal ion has been proposed to be involved in polymerase nucleotide catalysis and may play a role in these interactions (37–41). Interestingly, unlike Q151N, E710A had a larger effect on *k*_cat_ rather than on binding, leading the authors to postulate that its interaction with the 3′-OH does not contribute to ground state binding. Instead, the data was more consistent for E710-Mg^2+^ interactions having a role in the transition state occurring after nucleotide binding but before catalysis (36). Our work indicates that lowering the concentration of Mg^2+^ disproportionately depresses the rate of catalysis for all thymidine analogs lacking a 3′-OH without strongly affecting affinity (based on K_d_ values) for a limited set (dTTP, AZTTP, and d4TTP) of tested nucleotides (Fig. 3, and Tables 1 and 2). This is consistent with the observations with the KF and suggests that Mg^2+^ may interact with the 3′-OH, either directly or indirectly, in a step after nucleotide binding during the transition state or the chemistry step (the latter was ruled for the KF by demonstrating a lack of a sulfur elemental effect (36), but we did not test this possibility).

It was noteworthy that HIV RT was able to incorporate all 9 analogs that were tested. Only 3′-O-CH_3_ dTTP showed very poor incorporation. For chain terminating analogs, there was not a clear consistent chemical property that explained the different rates of incorporation of the analogs. However, electronegative groups that, unlike a 3′-OH, could not donate a H atom for hydrogen bonding (e.g. 3′-F and 3′-O-CH_3_) were incorporated the slowest. Groups with potential for positive charge or hydrogen bond formation (e.g. 3′-NH_2_, and 3′-N_3_), and small groups (e.g. 3′-H (ddTTP) or d4TTP (no group)) were better substrates, although all were less efficiently incorporated than dTTP, especially in low Mg^2+^ (Tables 1 and 2). It is possible that 3′-NH_2_ and 3′-N_3_ groups may interact with Q151 through charge interactions or H-bonding and this may help compensate for loss of the 3′-OH. Both ddTTP and d4TTP could not establish these interactions but the small size of the 3′ substitution and non-charge nature could minimize repulsive interactions. Also, d4TTP is different than the other tested chain terminators as the 2′,3′-didehydro configuration of the sugar moiety alters the structure of the sugar. This makes it more difficult to directly compare it to the other 3′ modified analogs.

In conclusion, our results support a role for interactions between Mg^2+^ and the 3′-OH of the incoming nucleotide that enhances the rate of catalysis. Interactions between Mg^2+^ and the 3′-OH terminus of the growing DNA chain and the incoming nucleotide phosphate backbone have been well established and are part of polymerase dogma. In these instances, divalent cation is directly interacting with the reacting moieties of nucleotide catalysis. Presumably, the 3′-OH of the incoming nucleotide is spatially removed from the catalytic center. This suggests that an interaction with it that enhances catalysis would likely have a stabilizing (e.g. stabilization of a transition state) rather than chemical role. An example could be that Mg^2+^ effects established interactions between Q151 and the 3′-OH. This could occur directly or indirectly through positioning of the nucleotide in a manner that promotes the Q151/3′-OH interaction. Further experiments, including those with Q151 mutants and mutations at other positions that effect Mg^2+^ binding, will be required to better understand the mechanism behind these observations.

## Materials and Methods

### Materials

3′-fluor (F) and 3′-O-methyl (O-CH_3_) dTTP were synthesized by AX Molecules Inc., San Marcos, CA. 3′-amino (NH_2_) dTTP and dideoxy TTP (ddTTP) were from TriLink Bio Technologies. AZTTP and d4TTP were from Sierra Bioresearch, Tucson, AZ. 4′-C-methyl, 4′-C-ethyl, and D-carba dTTP (23, 24) were a generous gift from Dr. Stephen Hughes (National Institutes of Health). All other nucleotides were from Roche Diagnostics Corporation. All nucleotides had purity levels >95% (as determined by mass spectroscopy and TLC analysis). T4 polynucleotide kinase (PNK) was from New England Biolabs. Radiolabeled compounds were from PerkinElmer. DNA oligonucleotides were from Integrated DNA Technologies (IDT). G-25 spin columns were from Harvard Apparatus. All other chemicals were obtained from Fisher Scientific, VWR, or Sigma.

### Preparation of HIV reverse transcriptase (HIV RT)

The expression plasmids (pRT66 and pRT51) for HIV-1 RT (HXB2 sequence) were used to produce HIV RT (25). The enzyme, which is a non-tagged heterodimer consisting of equal proportions of p66 and p51, was prepared as described (26).

### Preparation of primer-templates for pre-steady-state kinetics and inhibitor analysis on the 100-nt template

For pre-steady-state kinetics, 5′-^32^P-labeled 20-nt primer (5′-TTCCCGTCAGCCTAGCTGAG-3′) and 36-nt template (5′-ATCGTCGTACTCAGCTAGGCTGACGGGAAATTGTAT-3′) were mixed at a ratio of 1:1.25 (primer:template) in a buffer containing 50 mM Tris-HCl (pH 8), 80 mM KCl, and 1 mM DTT. For inhibitor analysis during primer extension on a long template, 5′-^32^P-labeled 20-nt primer (5′-TTGTTGTCTCTTCCCCAAAC-3′) and 100-nt template (5′-TGGCCTTCCCACAAGGGAAGGCCAGGGAATTTTCTTCAGAGCAGACCAGAGCCAACAGCCCCACCAGAAGAGAGCTTCAGGTTTGGGGAAGAGACAACAA-3′) were mixed at a at a ratio of 1:1.5 (primer:template). Hybrids were formed by heating to 80◻C for 5 min then slow cooling in a PCR machine to 4□C.

### Pre-steady-state kinetic parameters for dTTP or dTTP analog incorporation

Experiments were performed using a QFM-4000 from Bio-Logic Scientific Instruments. All experiments were carried out at 37□C. For each reaction, 20 μL of solution containing 50 nM (in primer) primer-template prepared as described above (final concentration 25 nM in reactions) and 50 nM HIV-1 RT (HXB2) (final concentration 25 nM in reactions) in buffer 1 (50 mM Tris-HCl (pH 8), 80 mM KCl, 1 mM DTT, and 0.5 or 6 mM MgCl_2_) were loaded into one loop of the apparatus. A second loop contained 20 μL of a solution with various concentrations of dTTP or specific dTTP analogs in the same buffer. Concentrations used ranged from 1 to 100 μM (final concentrations in reaction) for experiments to determine K_d_ and *k*_cat_ (depending on the nucleotide) and were 5 μM for pre-steady-state rate (*k*) determinations with various analogs. For reactions with higher nucleotide concentrations, the starting Mg^2+^ concentration in the 0.5 mM Mg^2+^ reactions was adjusted to keep the free Mg^2+^ concentration at 0.5 mM (19). A third loop contained 20 μL of a solution containing 0.5 M EDTA (pH 8). Reactions were initiated by rapidly mixing the solutions in loops 1 and 2, then terminated by mixing with loop 3 at time points ranging from 5 milliseconds (ms) to 2 seconds (s). Specific time points depended on the incorporation rate of the individual analogs. The products were mixed with and equal volume of 2X loading buffer (90% formamide, 0.025% each bromophenol blue and xylene cyanol) and resolved on 20% polyacrylamide-7M urea denaturing PAGE gels as described (27). Gels were dried, then exposed to phosphorimager screens. Quantification was performed using a Fujifilm FLA7000 phosphorimager and Fuji ImageQuant software. First order rate constants for incorporation (*k*) were determined using SigmaPlot by plotting the concentration of extended starting material vs. time, and fitting the data to a single exponential equation (28):

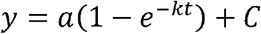

where “a” is the amplitude, *k* is the rate, and “C” is the end point. In some cases, especially for analogs with slower incorporation rates, the data fit better to a simpler equation:

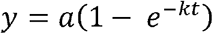

For some analogs, rate constants at different analog concentrations were used to determine the equilibrium dissociation constant and maximum rate of nucleotide addition (K_d_ and *k*_pol_, respectively) by plotting *k* vs. the concentration of nucleotide and fitting to a hyperbolic equation for ligand binding with single site saturation:

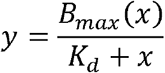

where “B_max_” corresponds to *k*_pol_. Standard error values for K_d_ and *k*_pol_ were generated by the program.

### Inhibitor analysis during primer extension on a long template

Primer extension reactions were performed to study the inhibition of extension by thymidine analogs as described previously (19). Briefly, 15 nM 5′-^32^P-labeled primer was hybridized with 22.5 nM template at a ratio of 1:1.5 as described above. Hybrids were preincubated for 3 min at 37°C in 8.5 μL of buffer 1, all 4 dNTPs and 1 of the analogs (5 μM each and 3 μM, respectively (final concentrations in reactions)). Extension was initiated by adding 4 μL of HIV RT (final concentration 100 nM) in buffer 1. After extension for 30 min, the reactions were terminated by adding 12.5 μL of 2X gel loading buffer and samples were resolved on an 8% denaturing urea gel and processed as described above.

